# Early Path Dominance as a Principle for Neurodevelopment

**DOI:** 10.1101/2022.07.14.500044

**Authors:** Rostam M Razban, Jonathan Asher Pachter, Ken A Dill, Lilianne R Mujica-Parodi

## Abstract

We perform *targeted attack*, a systematic computational unlinking of the network, to analyze its effects on global communication across the network through its *giant cluster*. Across diffusion magnetic resonance images from individuals in the UK Biobank, Adolescent Brain Cognitive Development Study and Developing Human Connectome Project, we find that targeted attack procedures on increasing white matter tract lengths and densities are remarkably invariant to aging and disease. Time-reversing the attack computation suggests a mechanism for how brains develop, for which we derive an analytical equation using percolation theory. Based on a close match between theory and experiment, our results demonstrate that tracts are limited to emanate from regions already in the giant cluster and tracts that appear earliest in neurodevelopment are those that become the longest and densest.

**Significance:** As brains develop through neural growth and specialization, what mechanism ensures that new neurons are integrated into a fully connected brain, avoiding “bridges to nowhere”? Here, we study brain structure development from the perspective of percolation, a global measure of communication. Analyzing over 35,000 diffusion MRI scans on human individuals, from newborns to adults, we identify the following rules of brain neurogenesis through percolation theory: earlier tracts become longer and denser while maintaining a giant cluster. This signature, invariant to age or mental health, suggests a fundamental condition for the brain to function as an emergent whole.

---

Brains are networks of neuronal regions (nodes) that are linked together by bundles of axons (edges). Much more is known about *static topologies* of brain networks than of the *dynamics* of how topologies emerge through neurodevelopment. Known topological properties include their degree distributions, path lengths, clustering coefficients and rich-club coefficients [1, 2, 3, 4, 5, 6, 7, 8, 9]. These graph-theoretic features can be captured by modeling, and compared to other known types of networks. For example, the high average clustering coefficients and low average path lengths in brains are similar to what is found in a lattice of sites with introduced random connections, called small-world networks [1, 5].

However, these topological properties of brains are often considered as snapshots at a given time [10]. Our interest here is in the developmental trajectories through which brains come to have its unique topology. There are no publicly available data sets yet rich enough to give the full trajectories of topologies of developing brains^1^. Thus, as an alternative, we use data to construct a generative model of a quasi-dynamical sequence of events of developing brain networks. We do this using statistical mechanical percolation theory with a procedure called *targeted attack* [13, 14, 15].

## Targeted attack analysis of brain data

*Targeted attack* is a computational procedure that can be performed on any network having a known topology. Targeted attack entails sequential removal *in silico* of nodes or edges based on the rank order of some node or edge property [13, 16, 14], followed by analysis of how the network changes step-by-step throughout the decimation of the net by the attack. Consider the metaphor of a city’s roads. One metric is road lengths: you might remove them in order of shortest to longest, for example. By a different metric, you might remove links in order of how many cities they connect. The nature of how a network topology diminishes throughout the attack can give insights about the network beyond what single network measures can give. In particular, we focus on the *giant cluster* (the largest group of connected nodes) because it is a proxy for global communication. We compute the probability *P* that an arbitrary node in the network is in the giant cluster. If *P* = 1, then any arbitrary region can communicate with any other region [17, 18].

Various targeted attack analyses have previously been performed on brain network data [2, 19, 20, 21, 22, 23, 24]. It has been found that when attack sequences are based on correlations of *functional connectivities* (e.g. functional MRI (fMRI) signals between regions), the brain does not decimate in the same way as random or scale-free graphs [2]. The attack procedure has also been applied to *structural connectivities* (e.g. diffusion MRI (dMRI)), which measures the topology of large bundles of axons known as *white matter tracts* [25]. This study removed nodes based on their degrees [26], motivated by the logic that their targeted attack procedure directly simulates neurodegeneration, which is more likely to be located in metabolically costly hubs [27].

Here, we seek new insights through the following novel aspects: (1) two new decimation strategies: tract lengths and densities; and (2) an analytical theory for time-reversing targeted attack, which can shed light on how the brain network builds up in this process. The logic is that time-reversal of the decimation process gives a plausible hypothesis for how brain structure emerges through early development, which we validate on *N*=35,731 human dMRI scans of increasing targeted attack of tract length and density acquired from the UK Biobank [28], the Adolescent Brain Cognitive Development (ABCD) Study [29] and the Developing Human Connectome Project (dHCP) [30]. We compare brain results to alternative graph theoretical structures, such as random, scale-free and small-world graphs, as well as a graph constrained by brain volume. And, to confirm that our conclusions are not specific to the dMRI data or human brains, we replicate our results on mouse viral tracing experiments [31].

## The computational procedure of targeted attack

At a given step of decimation, the various nodes of the remaining graph will have different degrees of connectivity (numbers of other nodes to which it connects). At that stage of decimation, the graph’s average connectivity will be ⟨*k*⟩. The calculation we make here is of *P* = *P* (⟨*k*⟩). For example, node c in Figure 1 initially has a degree (*k*) of 2 in the left-most realization of the graph “abcd.” ⟨*k*⟩ is the average over all nodes’ degrees. *P* is calculated by the number of nodes in the giant cluster divided by the total number of nodes. The left-most realization of the graph “abcd” first has *P* = 4*/*4, then sequential attack from small to large tract lengths results in *P* = 4*/*4, *P* = 3*/*4 and *P* = 2*/*4 (Figure 1).

**Figure 1:**
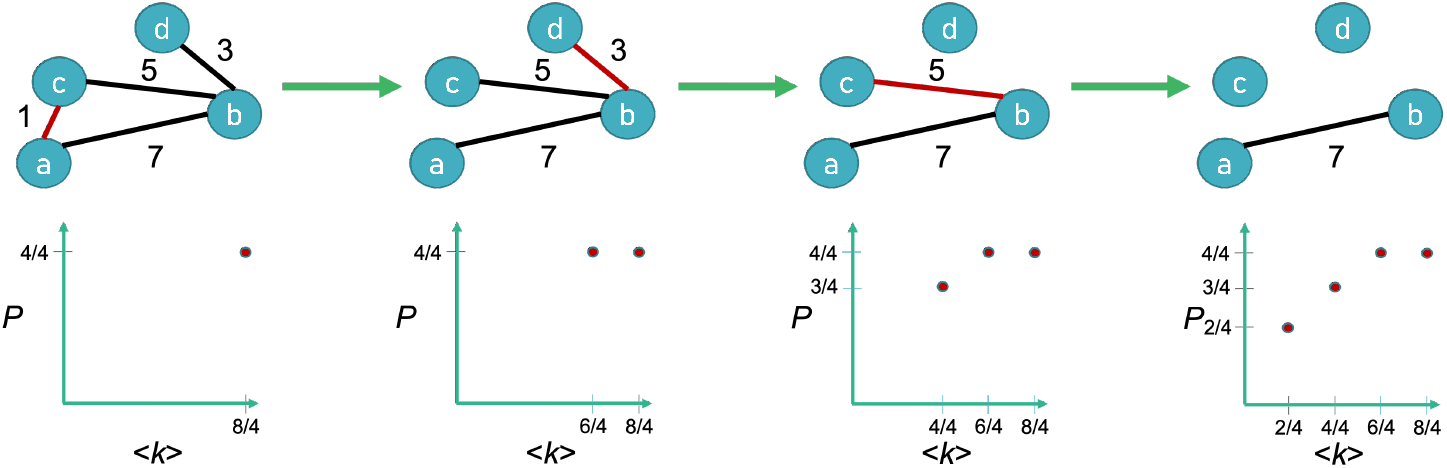
The targeted attack procedure on increasing tract lengths. By starting with all edges present on the left, we sequentially remove those edges whose length (listed above the edge) are currently the shortest (colored in red). *P* is the probability a node is in the giant cluster; ⟨*k*⟩ is the average degree.

We run a targeted attack procedure on two main outputs from a generic dMRI data analysis: *tract length* and *tract density* (Figure 1). Tract density, also known as streamline count, is the number of tracts connecting two grey matter regions; tract length is the average length of those connections (Methods). Since we can remove edges in *increasing* (smallest to largest) or *decreasing* (largest to smallest) order, each physical property has two possible *P* curves. Figure 2 demonstrates that removal of increasing tract lengths and tract densities yield *P* curves qualitatively different from those corresponding to random graphs. In contrast, decreasing tract density removal results in a poorly resolved curve because a high proportion of edges have a tract density of 1 (Figure S1, right). Decreasing tract length removal results in a *P* curve qualitatively similar to that of random graphs (Figure S1, left), consistent with previous results [26]. Here, we analyze the shapes of the curves of *P* (⟨*k*⟩) from increasing tract length and tract density targeted attack, and compare to other known types of networks.

**Figure 2:**
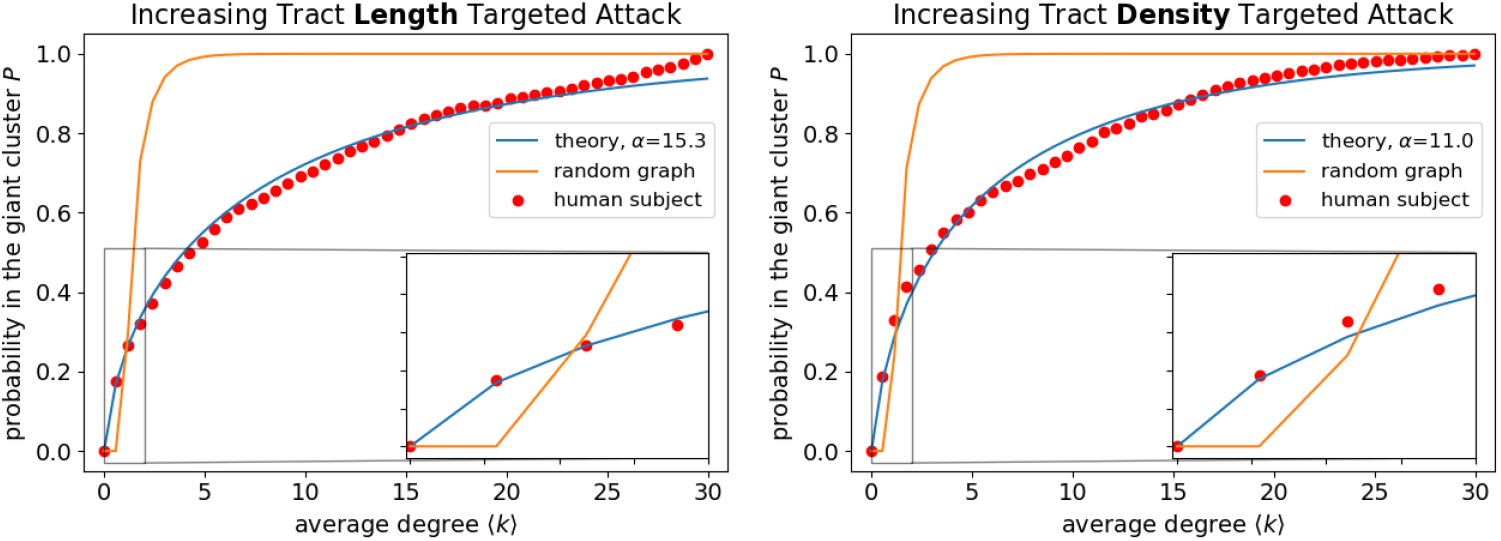
Attack curves, *P* (⟨*k*⟩), theory vs. experiments. (Red points) dMRI data collected by the UK Biobank for one arbitrarily chosen human individual (subject ID: 6025360) [33] under the Talairach atlas [34]. (Blue line) The presented Giant Cluster Self Preference theory. (Orange line) Prediction from random graph theory. The inset zooms in on the first four equidistant sampled experimental points to highlight the lack of a sharp transition.

## Analytical theory for brain network formation

The attack procedure describes the *P* curve from right to left, by a breakdown process. Instead, looking at the same *P* curve now from left to right, describes a hypothetical build-up process that provides a growth trajectory on network development from early to late. One well-known trajectory in percolation theory is *a random network*. Given in terms of the Lambert *W* function [15], an analytical solution is known for *P* (⟨*k*⟩) in the limit of infinite numbers of nodes [32, 15].

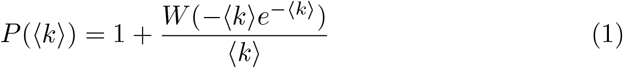

We are not aware of any other known networks for which *P* (⟨*k*⟩) is fully derived ^2^. Here, we analytically characterize a novel mechanism for growth of the giant cluster by restricting secondary cluster formation, based on experimental evidence discussed in the next section. Secondary clusters are defined as other clusters besides the giant cluster of size greater than one. We compute the trajectory *P* (⟨*k*⟩) of what we call the *Giant Cluster Self Preference network* over its full range of ⟨*k*⟩, as (derived in the Methods section):

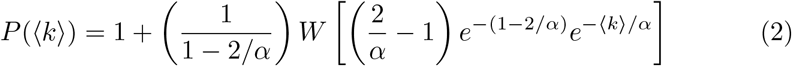

Equation 2 contains only one parameter, *α*, which we fit to the experimental data. The parameter reflects how many times more likely a new link lands in the giant cluster, relative to landing in a newly formed isolated node during the process of brain growth in early development.

## Results

### Attack trajectories depend on tract lengths and densities

Figure 2 demonstrates that for both tract length and tract density, random graph theory is not consistent with the experimental data. Experiments show that *P* (⟨*k*⟩) changes much more gradually all the way down to ⟨*k*⟩ = 0 at the left end of the figure. In one perspective, this might be surprising. Percolation behaviors often have steep cliffs: no global communication below a certain critical threshold, then a jump to a finite value of communication above it [14, 15]. These two curves are for one adult individual; Figures S2 shows that other adult brains have similar features.

We then probe more deeply into the difference of the brain network vs. random graphs. At ⟨*k*⟩ = 1, the giant component in the brain is essentially the only cluster; nodes not in the giant cluster are isolated and do not form secondary clusters. In contrast for the random graph at ⟨*k*⟩ = 1, the giant cluster is just one of many non-negligible clusters present (see Figure S3). This holds across different values of ⟨*k*⟩ (Figure S4). We conclude that in this particular computational attack, the brain’s giant cluster continuously degrades one node at a time, rather than by sharp fragmentation into secondary clusters from one large cluster. This lack of fragmentation explains the lack of a critical point in the *P* curves. We encode this mechanism into our Giant Cluster Self Preference theory (Equation 2) and find good agreement of the full attack curves *P* (⟨*k*⟩), for both tract length and tract density attack variables (Figure 2).

To rule out other potential mechanisms, we perform simulations of a network build up by preferential attachment [35]. We find a match with experimental *P* curves at a coarser parcellation (Figure S5). Scale-free networks built by preferential attachment are known to lack critical points during random attack [16, 14, 19]. However, the match fails to extend to finer parcellations (Figure S6) even when considering general preferential attachment models where edge addition is not simply linear in nodes’ degrees (Figures S7). The Giant Cluster Self Preference theory better captures trends across different parcellations (Figures 2 and S8).

Rather than building up graphs, we also create final networks and perform targeted attack to test whether corresponding *P* curves match those of real brains. Two models we consider are small-world networks [1] and random edges with distances based on the center of mass coordinates of regions in the parcellation. Both are limited to tract distance; they provide no information on tract lengths nor on tract densities. Neither model exhibits a match with the corresponding experimental data for increasing targeted attack on tract distance at either a coarse or fine parcellation (Figures S9 and S10). In the case of small-world networks, this is not unexpected because it is essentially a 2-dimensional ring lattice and lattices are known to have critical points [14, 15].

### *α* as a universal property of brains

The Giant Cluster Self Preference theory uses a fit parameter *α*, which measures the relative rates of a new network edge landing inside the giant cluster versus landing outside the giant cluster (Methods). Does *α* reflect some universal feature of brains, or depend on different brain states? We test whether *α* depends on the age or mental health status of adults based on the biometric properties in the UK Biobank data set. We find no strong systematic changes in *α* as a function of age (Figure 3)^3^. We also find that *α* values are indistinguishable by gender (Figure S11). In addition, no differences are found in *α* for individuals diagnosed with diabetes compared to healthy individuals (Figure S12) or for individuals diagnosed with bipolar disorder or depression compared to healthy individuals (Figure S13). The shape of the attack/trajectory curves, and the *α* values for each of them appear to be a relatively universal feature of human adult brains.

**Figure 3:**
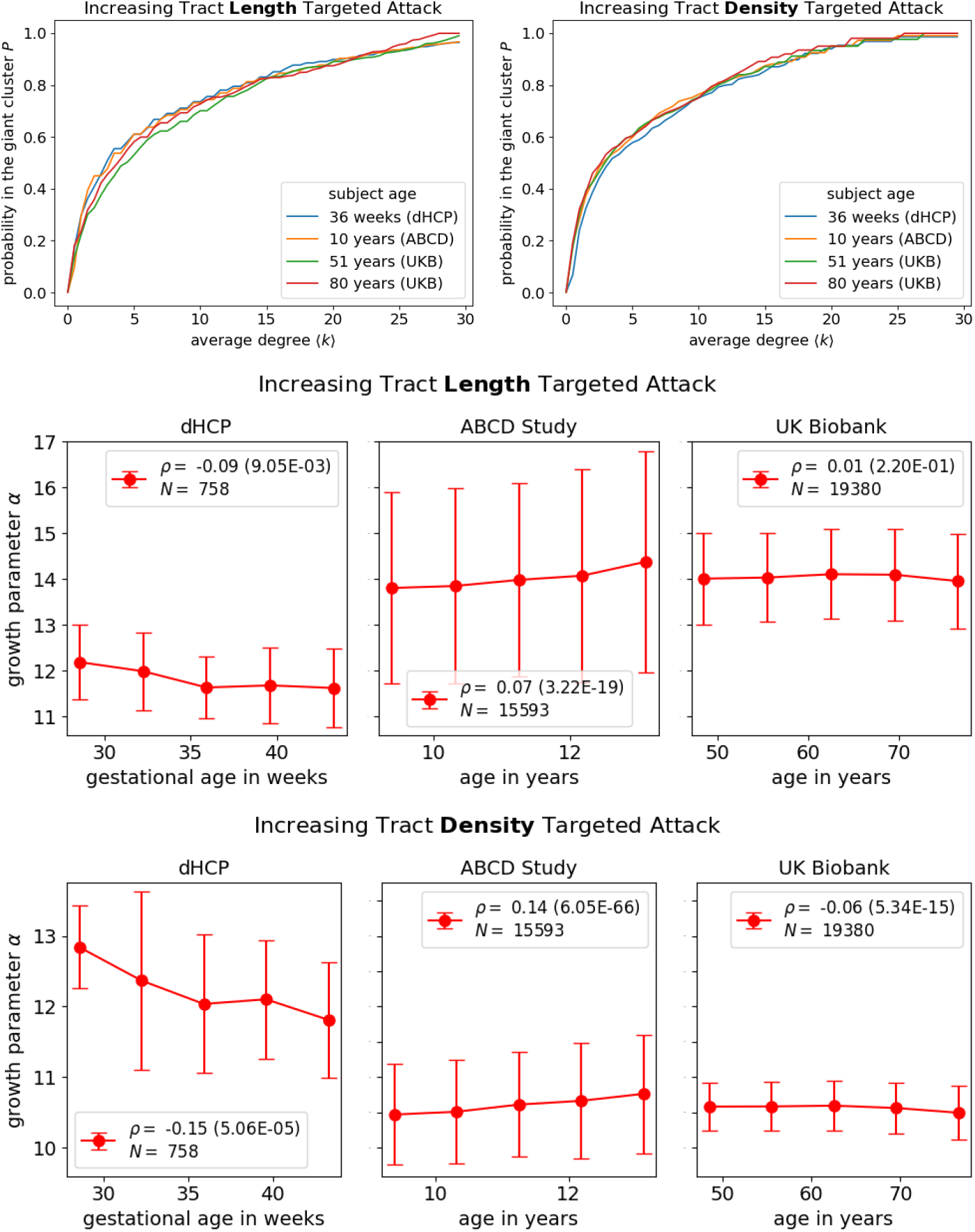
Attack curves for different human subjects of different ages. *P* (⟨*k*⟩) is remarkably similar across different ages within and across different data sets. To concisely capture the behavior of all *P* curves, we show fitted *α* parameters as a function of age in the bottom two rows of plots. Binned data are presented with a line connecting means, and error bars correspond to standard deviations. The variable *ρ* corresponds to the Spearman correlation coefficient calculated over all individuals between age and *α*, with p-value in parentheses. The variable *N* corresponds to the total number of individuals from the respective data set included in the analysis. An arbitrarily chosen sample of subjects are shown in the top row for subjects of age 36 weeks (Developing Human Connecto8me Project, subject ID: CC00063AN06, *α*_length_=12.7, *α*_density_=11.9), 10 years (Adolescent Brain Cognitive Development Study, subject ID: NDARINVNVF8N71U, *α*_length_=13.1, *α*_density_=10.8), 51 and 80 years old (UK Biobank, subject IDs: 6025360 and 4482035, *α*_length_=15.3, *α*_density_=11.0 and *α*_length_=14.2, *α*_density_=10.4, respectively).

To extend our results to younger subjects, we analyzed dMRI data from the ABCD Study [29], which contains individuals aged 9 through 13 years old. We find *α* values for the corresponding tract density and length targeted attack remarkably similar to those of adults studied in the UK Biobank and relatively insensitive to age, although more sensitive than those of the UK Biobank (Figure 3). Given the widespread presence of synaptic pruning during adolescence [36, 37, 38], our results indicate that synaptic pruning plays a minor role in affecting percolation because there are many edges responsible for maintaining *P* = 1. Furthermore, we analyze dMRI data from dHCP [30] for recently conceived newborns and reach the same conclusion, although the similarity with adult *α* values is not as high. Figure 3 illustrates the similarity in *P* curves across individuals of different ages. Taken together, these results imply that *α* does not reflect postnatal human white matter development, and is more consistent with prenatal development.

To check that our results are not an artifact of the dMRI data modality, we studied viral tracing data of mice from the Allen Institute [31]; see Figure S14. Again, the present Giant Cluster Self Preference theory gives excellent fits to the attack curves, but now with larger-than-human values of *α* for distance-based attacks and smaller-than-human for density-based attacks^4^.

Our claim that such a targeted attack approach can provide insight into prenatal development is not without precedent. It has direct support at the neuronal scale of mice [39, 40, 41]^5^. Taken together, our results across humans of different ages and mental health diagnoses, as well as on mice using a different data modality, support the claim that reversing the respective targeted attack procedure provides a hypothesis for how brains develop.

### During neurodevelopment, the earliest tracts become the densest and longest

One observation that the Giant Cluster Self Preference theory does not address is that targeted attack yields nonrandom *P* curves when attacking edges based on *increasing* tract lengths and densities. Results from the previous subsection indicate that the *P* curves reflect a special tract formation order during fetal development in which the brain is rapidly growing [42, 36]. In Figure 4A, we propose a model, *Early Path Dominance*, that supplements the Giant Cluster Self Preference theory by accounting for concurrent brain growth as the giant cluster increases in size. As the fetal brain is emerging, initial edges are short with small tract densities; however, as the brain grows in size and new regions appear, those same edges lengthen and become more dense. Tract lengthening has been called *scaling* and found to be present in the nervous system development of *C. elegans* [43, 44, 45, 9]. In the context of dMRI, there are multiple reasons that tracts can appear more dense. These include both *fasciculation* (increasing fiber count) [46, 42, 47], as well increasing myelination [25].

**Figure 4:**
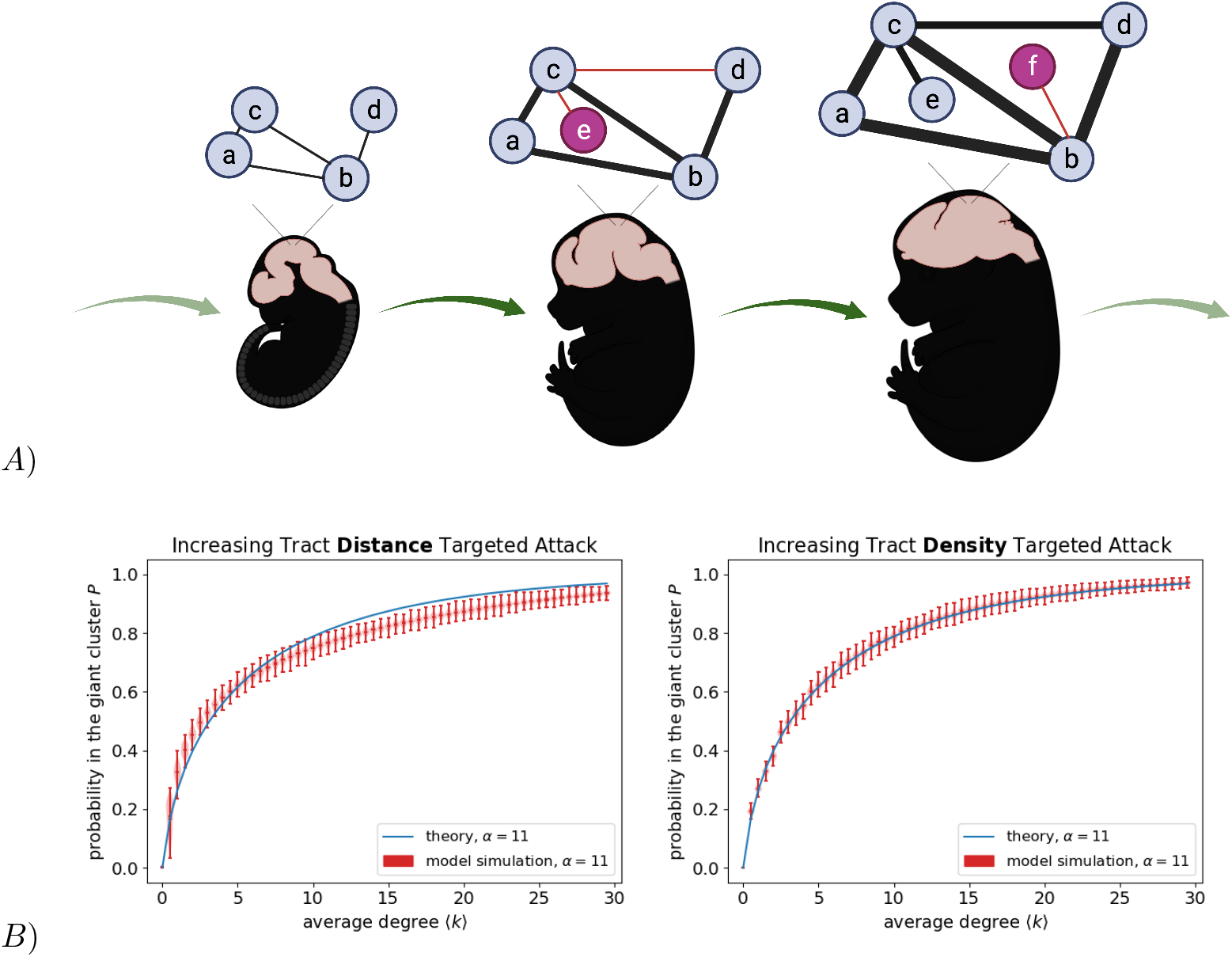
Early Path Dominance Model: Proposed sequence of topological growth in brain development. (A) New edges, marked in red, start off short and thin (less dense) and become longer and wider (more dense) with each growth step. Figure created with biorender.com. (B) Simulations of the model are consistent with theory for the same *α* parameter. Violin plots represent 1000 independent runs of a graph with 727 nodes, the same number as in the Talairach atlas.

We perform a simulation of the Early Path Dominance model, incorporating scaling and fasciculation/myelination alongside the critical constraint that edges only add between regions where at least one region is already in the giant cluster (Giant Cluster Self Preference theory). Physical growth is encoded by randomly assigning coordinates to nodes within a unit sphere and assuming existing coordinates uniformly increase by a fixed constant every time an edge is added (Methods). Figure 4B demonstrates that the *P* curves from the Early Path Dominance model agree well with the Giant Cluster Self Preference theory, which itself showed good agreement with experiments (Figure 2). The Giant Cluster Self Preference theory (Equation 2) makes no specific reference to tract length and density in so much that edge addition (increasing ⟨*k*⟩) reflects the corresponding tract order formation (further discussed in Methods).

An important feature seen in brain data is higher average clustering coefficients but similar average path lengths to random graph networks [48, 6, 49, 50]. The Early Path Dominance model captures this feature to a certain extent (Figure S16), despite not being specifically designed to do so ^6^. Small-world networks are capable of achieving the higher clustering coefficients seen in brain data at finer resolutions [1]. For the same adult individual as in Figure 2 under the Talairach atlas, we find that the average clustering coefficient is 0.60, while a random graph with the same average degree of 29.9 has an average clustering coefficient of 0.04. However, it is unclear whether highly elevated average clustering coefficients at fine parcellations are artifacts of the edge sparsity [51, 52].

### Wider Perspectives

The relative invariance of the *P* curves implicates that this percolation mechanism may matter for brain function, and that it may be a relatively universal mechanism of brain growth. Moreover, fitting brain data require values of *α >* 1, meaning new neurons have greater propensity to connect inside the giant cluster than outside it. Taken together, these points lead us to the following speculations. First, perhaps the preference of neurons to link to the giant cluster is because of a “Hebbian principle” [53], where neurons have greater recruitment and growth into regions of higher neuronal activity, which, in this case, may be associated with larger clusters. Future work will test this speculation with activity data of neuronal regions from functional MRI. Second, this mechanism has the efficiency advantage to the organism of “no neuron left behind”. That is, neurons mostly link to clusters that are active and connected. And third, this mechanism is opportunistic, generating new connections stochastically that lead to different brain wirings in detail, even while constrained by a common principle.

## Conclusions and Discussion

We have analyzed human brain network topologies by performing an attack analysis, systematically removing links in the computer and watching how it changes the probability of being in the giant cluster, *P* (⟨*k*⟩) as a function of the average degree ⟨*k*⟩ of the network at that stage. We use two different attack variables: tract length and density. These curve shapes under increasing targeted attack are universal across postnatal age and disease, and in mice, but are very different than for random networks. We hypothesize that time-reversing the attack procedure may mimic the physical neuronal development of fetal brains. On that basis, we derive an analytical equation that grows the brain network and provides two fundamental insights into white matter tract development. First, tracts form primarily from regions already in the giant cluster. Second, on average the first tracts constructed become the longest and the densest.

## Methods

### Calculating the connectivity matrix

Connectivity matrices are calculated using the Diffusion Imaging in Python (DIPY) software [54]. Alongside the already preprocessed dMRI images from the respective data set (discussed in the SI), we input a brain atlas that distinguishes between white and grey matter, as well as parcellates the grey matter into an arbitrary number of regions. In the Results, we use the Talairach atlas [34], however, we also show results for the Harvard-Oxford [55] and the modified Desikan-Killiany atlases [54, 56] in the Supplement (Figure S8). Nodes found to form zero edges when calculating the connectivity matrix are removed from consideration such that *P* = 1 when all edges are considered.

We perform a deterministic tracking method in DIPY to generate the connectivity matrix [54]. Reconstruction of the orientation distribution function is done using Constant Solid Angle (Q-Ball) with a spherical harmonic order of 6 [57]. The relative peak threshold is set to 0.8 with a minimum separation angle of 45 degrees. We only seed voxels in the white matter and count tracts that ended in the grey matter. Minimum step size of tracts is 0.5 mm. Our protocol to obtain connectivity matrices closely follows a DIPY tutorial found on their website under streamline tools. We tried a different parameter (spherical harmonic order = 8) and reconstruction method (Diffusion Tensor Imaging) to calculate connectivity matrices and found similar *P* curves as those shown in the main text (Figure S17). Results are also robust to variations in number of nodes or average degrees seen across individuals’ brain networks (Figure S18).

Tractography outputs the number of tracts that connects two grey matter regions, which we call the tract density, and the individual lengths of each tract. Because each tract can have its own unique length, we take the average over all lengths to represent a given connection between two grey matter regions and call that the tract length.

### The Giant Cluster Self Preference Theory

We seek a network with specific characteristics which quantitatively displays agreement with the wealth of percolation curves generated from targeted attack on dMRI data. In this section, we derive an analytical expression for *P* (⟨*k*⟩) - the fraction of nodes residing in the giant cluster as a function of the average degree ⟨*k*⟩ - for this network.

In the first subsection of the results, we observed that brain networks under increasing tract length and density targeted attack display the peculiar feature of essentially never having secondary clusters; there is just the one giant cluster, and then either isolated nodes (nodes that form no other edges) or very small clusters (Figures S3 and S4). To develop our analytical theory, we take this to the extreme and approximate brain networks as having only one cluster. For a growing brain network, we even expect no isolates. When performing the targeted attack in order to generate percolation curves, sometimes removing a link causes a node to become isolated, like an island that has been set free from a mainland, and those are the isolates we see in the deconstructed network. In our theory, we consider the reverse of targeted attack, i.e. the growth and development of the network over time. From this perspective, there is always just one cluster, and the appearance of a new node coincides with a new link that emerges from a random node in the cluster (see nodes *e* and *f* in Figure 4A).

Now consider a growing brain network currently containing *n* nodes, all in one cluster, with *E* edges. Each growth step corresponds to the addition of exactly one new edge - the only question is, will this new edge reside in the existing cluster, providing more direct connections for the nodes already there, or will it branch outward into a new node? Here we introduce the factors defining our analytical theory of brain networks.

There are *n*(*n* − 1)*/*2 − *E* available edge spots within the cluster. If a new node forms with some probability 1/*α*, then there are *N* − *n* possible new nodes that can connect to any of the *n* nodes in the cluster. *N* corresponds to the final number of nodes in the full-sized network. The *α* parameter captures brain growth with respect to parcellation. Although growth rates change throughout early development [58], *α* is simply taken to be a constant and fitted to the data^7^. An alternative yet equivalent interpretation is that an added edge is *α* times more likely to show up in one of the available spots within the giant cluster versus those connecting to a new region.

With the complete number of possible edges tabulated, we can calculate the probability *p*(*n* → *n* + 1 | *E* → *E* + 1) that we add a new node to the cluster during this growth step:

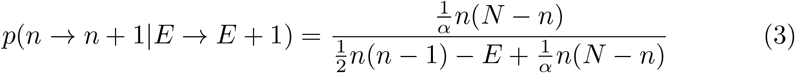

Since there are only two options at each growth step, the probability to place the new edge within the existing cluster rather than creating a new node is simply *p*(*n* → *n*|*E* → *E* + 1) = 1 − *p*(*n* → *n* + 1|*E* → *E* + 1).

We can now use these expressions for the transition probabilities to study how the probability *p*(*n* | *E*) of having *n* nodes evolves as more edges are added to grow the network:

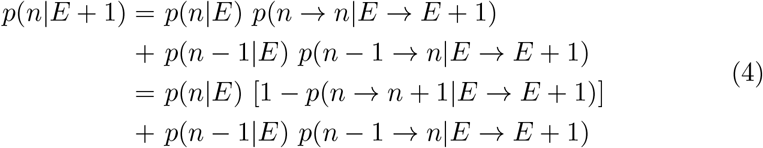

Rearranging slightly and plugging in the expression from Eq. (3) yields a discrete master equation:

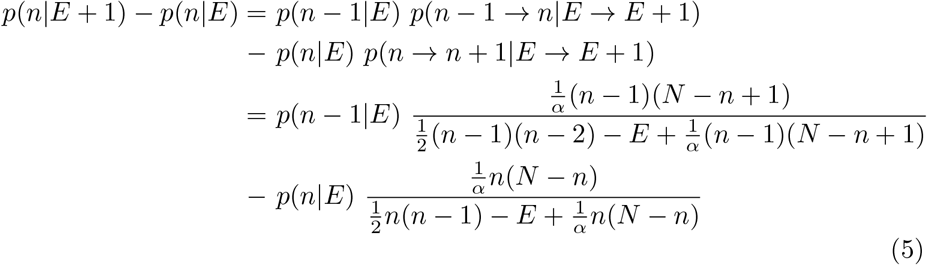

This difference equation can be solved numerically for any value of *N* to give the full evolution of *p*(*n*|*E*), using the initial condition *p*(*n*|0) = *δ*_*n*=1_. The function *δ* corresponds to the Kronecker delta; *δ*_*n*=1_ takes a value of 1 when *n* = 1 and is 0 elsewhere.

We now consider the limit as *N* → ∞, where both *n* and *E* also go to infinity but the ratios *ρ* = *n/N* and *κ* = 2*E/N* remain finite^8^. In this limit, defining *p*(*ρ*|*κ*) = lim_*N*→∞_ *Np*(*n* = *ρN* |*E* = *κN/*2) we can rewrite Eq. (5):

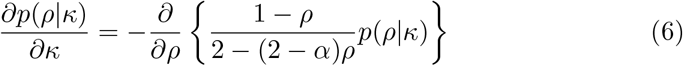

With initial condition *p*(*ρ*|0) = *δ*(*ρ*), the full solution of this partial differential equations is (using, for example, the method of characteristics) *p*(*ρ*|*κ*) = *δ*[*ρ* − *f* (*κ*)], where the function *f* (*κ*) satisfies:

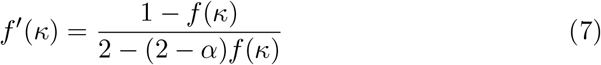

and *f* (0) = 0. The solution of this now ordinary differential equation is:

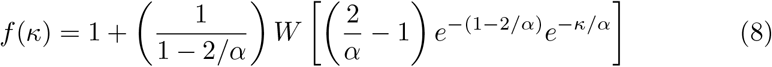

In this limit, the stochasticity in *p*(*ρ*|*κ*) is lost and the network size actually grows deterministically with growing edge density ratio *κ*, satisfying *ρ* = *f* (*κ*). Recognizing that *κ* - the edge density ratio in the growth perspective - corresponds to the average degree *k* in the targeted attack perspective^9^, and that *ρ*(*κ*) - the fractional size of the network compared to its full size as a function of *κ* in the growth perspective - corresponds to the probability *P* (⟨*k*⟩) to be in the giant cluster as a function of ⟨*k*⟩ in the targeted attack perspective, we arrive at Eq. (2), which we reproduce here for ease of reading:

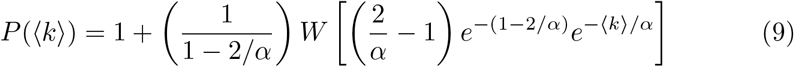

Although this form bears some resemblance to the analogous expression for a random graph (Eq. (1)), including the appearance of the Lambert *W* function, it produces significantly different features, particularly *P* (⟨*k*⟩) approaching 1 much more gradually with increasing ⟨*k*⟩. Eq. (9) matches the numerical solution of Eq. (5), validating our large *N* approximations (Figure S19).

This form matches targeted attack data on increasing tract density and length remarkably well, when fitted with an optimal *α* value (Figure 2). This further validates the main assumptions: brain networks have a single connected cluster throughout development, and internal connections are added continuously during growth, whereas the network only branches out and creates new nodes every so often. These aspects constitute our theory of developing brain networks, which we call Giant Cluster Self Preference. The name encapsulates the central idea that internal connections are shored up before the cluster branches out and connects with new nodes.

One limitation is that the Giant Cluster Self Preference theory does not include any reference to tract length or tract density. Future work will extend the current theory by assigning weights to each edge corresponding to these quantities and explain differences in fitted *α* values between tract density and length-derived *P* curves. Tract length and density are known to anticorrelate [59, 6, 60] (Figure S20). However the anticorrelation does not underlie results because increasing tract length *and* tract density targeted attack produce qualitatively similar results rather than length/density increasing and density/length decreasing (Figure 2 and S1). Furthermore, future work could build on the foundations here to explore other graph-theoretic aspects of networks governed by the Giant Cluster Self Preference theory, such as degree distributions, and how these quantities evolve over time.

### Simulating the Early Path Dominance Model

In Figure 4B, we demonstrate that corresponding *P* curves for simulations of the Early Path Dominance model matches the Giant Cluster Self Preference theory and thus, are consistent with experimental *P* curves (Figure 2). In principle, there are numerous ways of implementing the Early Path Dominance model according to the rules outlined in Figure 4A. Here, we constrain nodes to originate randomly within a sphere centered at the origin of radius 1. Nodes’ Cartesian coordinates are rescaled by 1.0001 each time an edge is added to the network, causing existing edges to lengthen by a factor of 1.0001. New edges are initially given a density of 1 that increases by a factor of 1.001 each time an edge is added to the network, causing existing edges to become more dense. Edges are added according to the Giant Cluster Self Preference theory with *α* = 11. Simulations are run with 727 nodes, the same number as in the Talairach atlas, and a final average degree of 100 to ensure that at the conclusion of the simulation, all graphs have *P* =1. Note that the additional randomness of coordinate placement seems to result in the distance targeted attack *P* curve requiring a higher *α* value (*α* ≈ 13) to capture the simulation (Figure 4B).

## Code and data availability

Scripts necessary to reproduce figures and conclusions reached in the text can be found at github.com/rrazban/percolating brain. Please refer to the respective publicly available diffusion MRI data set to access images.

## Acknowledgments

The authors thank Richard Granger, Elisha Moses, Botond Antal, Charles Kocher and Mobolaji Williams for helpful discussions of the results, as well as Botond Antal and Syed Fahad Sultan for assistance with the dMRI data. The research described in this paper is funded by the W.M. Keck Foundation (to LRMP and KAD) and the White House Brain Research Through Advancing Innovative Technologies (BRAIN) Initiative (NSFNCS-FR 1926781 to LRMP and KAD), and the Stony Brook University Laufer Center for Physical and Quantitative Biology (KAD).

This research has been conducted using the UK Biobank Resource under Application Number 37462.

Data were provided by the developing Human Connectome Project, KCL-Imperial-Oxford Consortium funded by the European Research Council under the European Union Seventh Framework Programme (FP/2007-2013) / ERC Grant Agreement no. [319456]. We are grateful to the families who generously supported this trial.

Data used in the preparation of this article were obtained from the Adolescent Brain Cognitive Development^SM^ (ABCD) Study (https://abcdstudy.org), held in the NIMH Data Archive (NDA). This is a multisite, longitudinal study designed to recruit more than 10,000 children age 9-10 and follow them over 10 years into early adulthood. The ABCD Study® is supported by the National Institutes of Health and additional federal partners under award numbers U01DA041048, U01DA050989, U01DA051016, U01DA041022, U01DA051018, U01DA051037, U01DA050987, U01DA041174, U01DA041106, U01DA041117, U01DA041028, U01DA041134, U01DA050988, U01DA051039, U01DA041156, U01DA041025, U01DA041120, U01DA051038, U01DA041148, U01DA041093, U01DA041089, U24DA041123, U24DA041147. A full list of supporters is available at https://abcdstudy.org/federal-partners.html. A listing of participating sites and a complete listing of the study investigators can be found at https://abcdstudy.org/consortium_members/. ABCD consortium investigators designed and implemented the study and/or provided data but did not necessarily participate in the analysis or writing of this report. This manuscript reflects the views of the authors and may not reflect the opinions or views of the NIH or ABCD consortium investigators. The ABCD data repository grows and changes over time. The ABCD data used in this report came from [NIMH Data Archive Digital Object Identifier 10.15154/1526472].

## Author Contributions

R.M.R., J.A.P. K.A.D., L.R.M.P. designed research, analyzed data, and wrote the manuscript. R.M.R. and J.A.P. performed the research.

## Supplementary Information

### Diffusion MRI data sets

#### United Kingdom Biobank

The United Kingdom (UK) Biobank is an ongoing epidemiological study aimed at discovering early markers for brain diseases [28]. UK Biobank is an excellent data source because of its large sample size (current release at our disposal: 19,380 people, ages 45 to 80 (mean age is 62.6 ± 7.4 years old)) and data standardization procedures – all corresponding measurements are collected on the same machines across several sites operating the same software [33]. More details on the minimally preprocessed dMRI images publicly assessed from UK Biobank can be found in Alfaro-Almagro et al. [33].

Individuals’ brain networks under the Talairach atlas have 727.4 ± 2.6 nodes with an average degree of 28.3 ± 1.1 (prior to targeted attack). The Spearman correlation between tract length and density is −0.20 ± 0.02 (Figure S20).

#### Adolescent Brain Cognitive Development Study

The Adolescent Brain Cognitive Development (ABCD) Study is an ongoing longitudinal study of ten years focused on understanding brain development during the critical period of adolescence [29]. The ABCD Study complements the UK Biobank by having individuals with an age range of 8.9 to 13.3 (mean age is 10.6 ± 1.1 years old), for the current data release at our disposal of 15,593 people (release 4.0, NIMH Data Archive Digital Object Identifier 10.15154/1526472). More details on the minimally preprocessed dMRI images of the ABCD Study publicly assessed from the National Institute of Mental Health Data Archive can be found in Hagler et al. [61].

Individuals’ brain networks under the Talairach atlas have 731.3 ± 2.6 nodes with an average degree of 31.0 ± 3.5 (prior to targeted attack). The Spearman correlation between tract length and density is −0.28 ± 0.05 (Figure S20).

#### Developing Human Connectome Project

The Developing Human Connectome Project (dHCP) acquires MRI from neonates within weeks of birth to probe early development [30]. The dHCP contains the youngest human subjects publicly accessible, gestational ages 26.7 to 45.1 weeks old (mean gestational age 39.7 ± 3.5 weeks old). We have 758 scans at our disposal from the current data release (third data release). More details on the minimally preprocessed dMRI images publicly assessed from dHCP can be found in Christiaens et al. [62].

Individuals’ brain networks under the Talairach atlas have 733.4 ± 1.9 nodes with an average degree of 34.4 ± 2.6 (prior to targeted attack). The Spearman correlation between tract length and density is −0.27 ± 0.03 (Figure S20).

**Figure S1:**
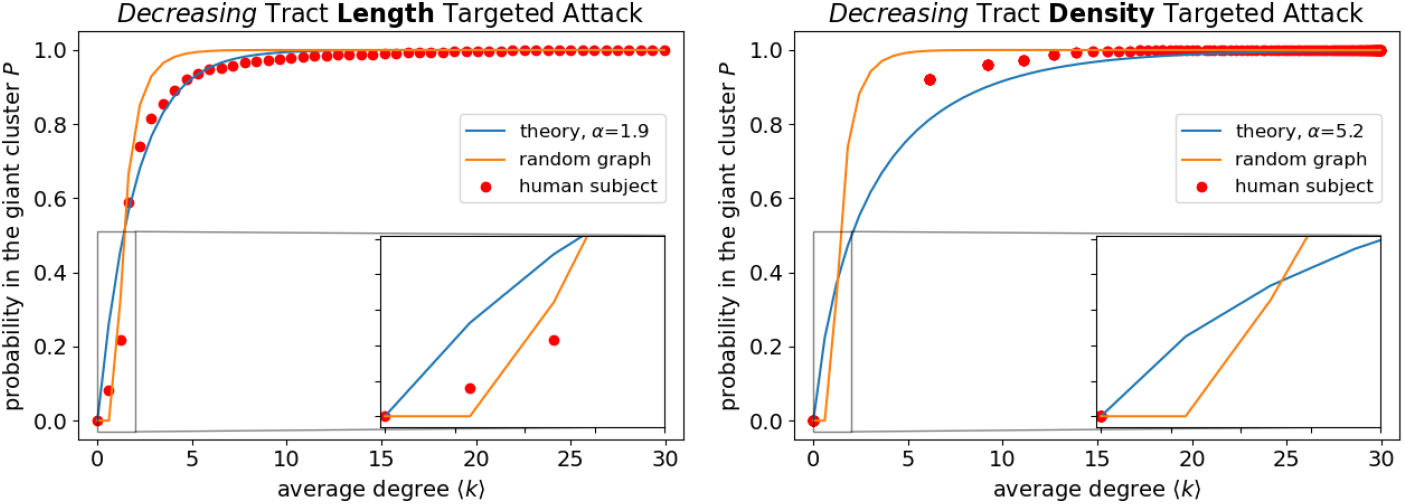
Targeted attack on *decreasing* tract lengths and tract densities. Note that throughout the main text, we focus on targeted attack of increasing tract lengths and densities and derive an analytical solution based on the Giant Cluster Self Preference theory (Equation 2), which is also shown in this figure for comparison. Targeted attack on decreasing tract lengths essentially follows a random graph. It is not possible to fully assess the corresponding *P* curve of tract density because of the large gap at low average degree. Curves are derived from dMRI data collected by the UK Biobank under the Talairach atlas for one human individual (subject ID: 6025360), the same subject as in Figure 2.

**Figure S2:**
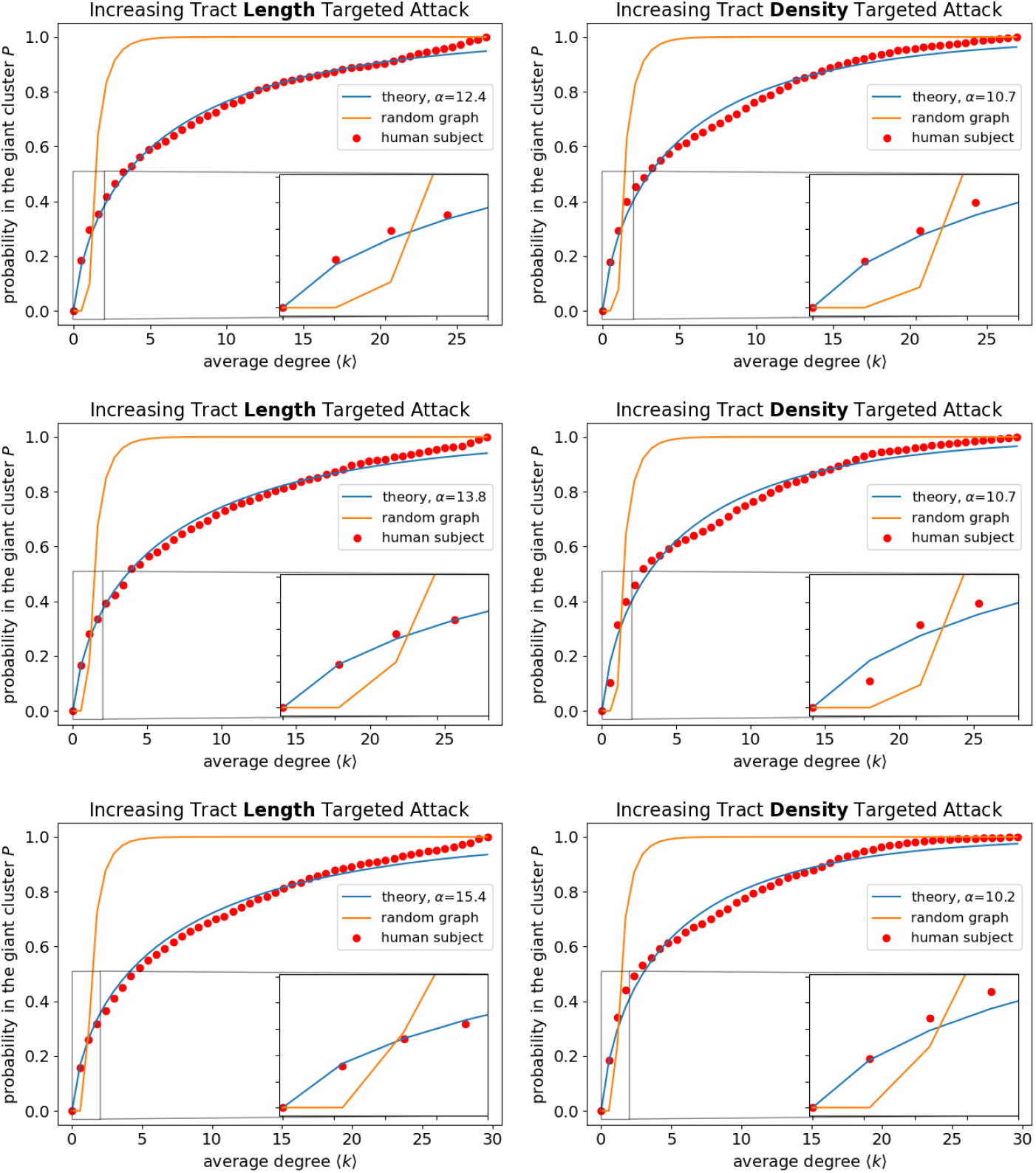
*P* curves for additional individuals besides that shown in Figure 2. From the top to bottom row, the UK Biobank subject IDs are 1701210, 1363111 and 2460658. The fitted *α* parameters are fairly similar among different subjects.

**Figure S3:**
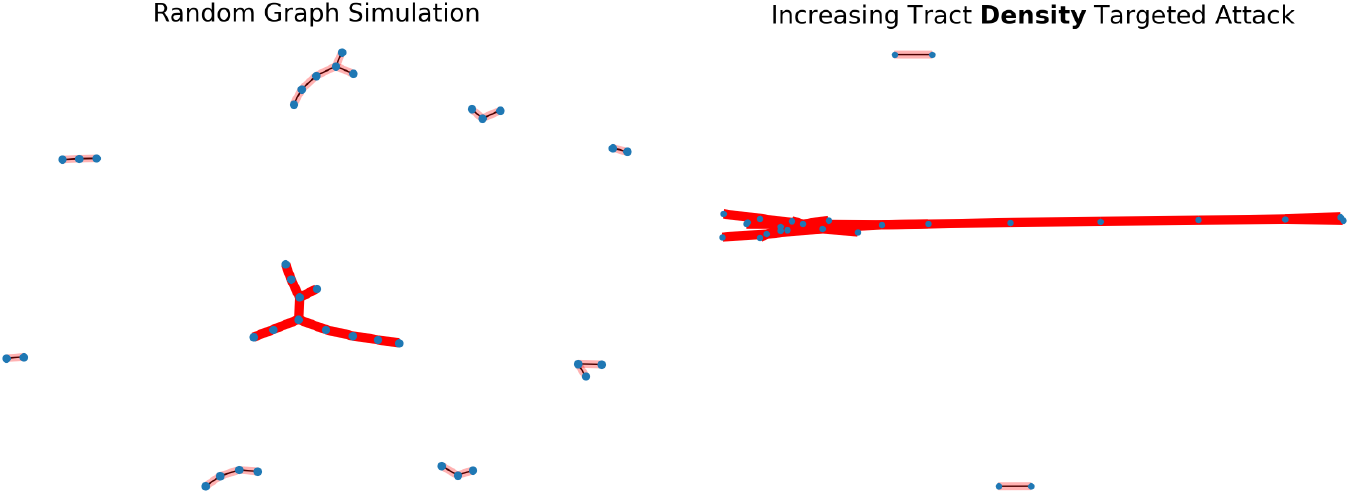
Visualizing cluster formation at average degree ⟨*k*⟩ ∼ 1 for both a random graph simulation (left) and the brain under increasing tract density targeted attack (right). The random graph simulation has several non-negligible secondary clusters, while the brain essentially has only one densely connected, large cluster. The random graph simulation has the same number of nodes as that of the brain shown analyzed under the Harvard-Oxford atlas, 64 total regions, however we exclude isolated nodes in both graphs for ease of visualization. The brain’s cluster distribution is derived from dMRI data collected by the UK Biobank for one human individual (subject ID: 6025360), the same subject as in Figure 2, however, analyzed at a coarser parcellation to easily visualize clusters.

**Figure S4:**
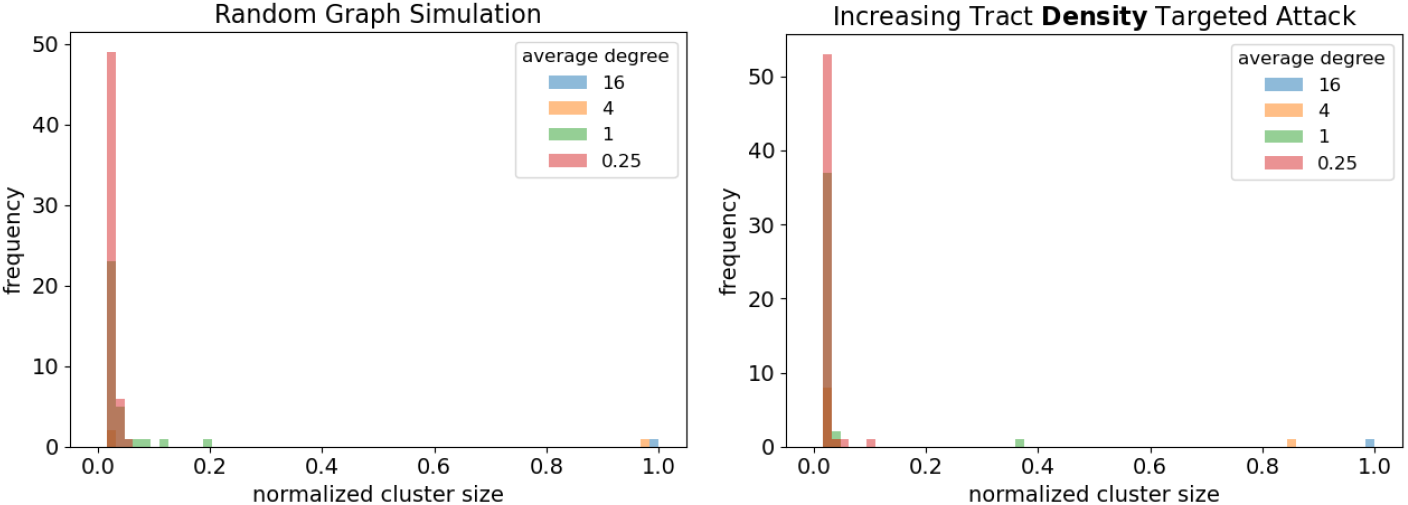
Cluster distributions across several average degrees for both a random graph simulation (left) and the brain under increasing tract density targeted attack (right). The random graph simulation has several non-negligible secondary clusters at small ⟨*k*⟩, while the brain essentially has only one densely connected, large cluster throughout. The random graph simulation has the same number of nodes as that of the brain shown analyzed under the Harvard-Oxford atlas, 64 total regions, as in Figure S3. Normalized cluster size is equal to cluster size divided by 64. Frequency refers to the number of different nodes with that cluster size. The brain’s cluster distribution is derived from dMRI data collected by the UK Biobank for one human individual (subject ID: 6025360), the same subject as in Figure 2, however, analyzed at a coarser parcellation.

**Figure S5:**
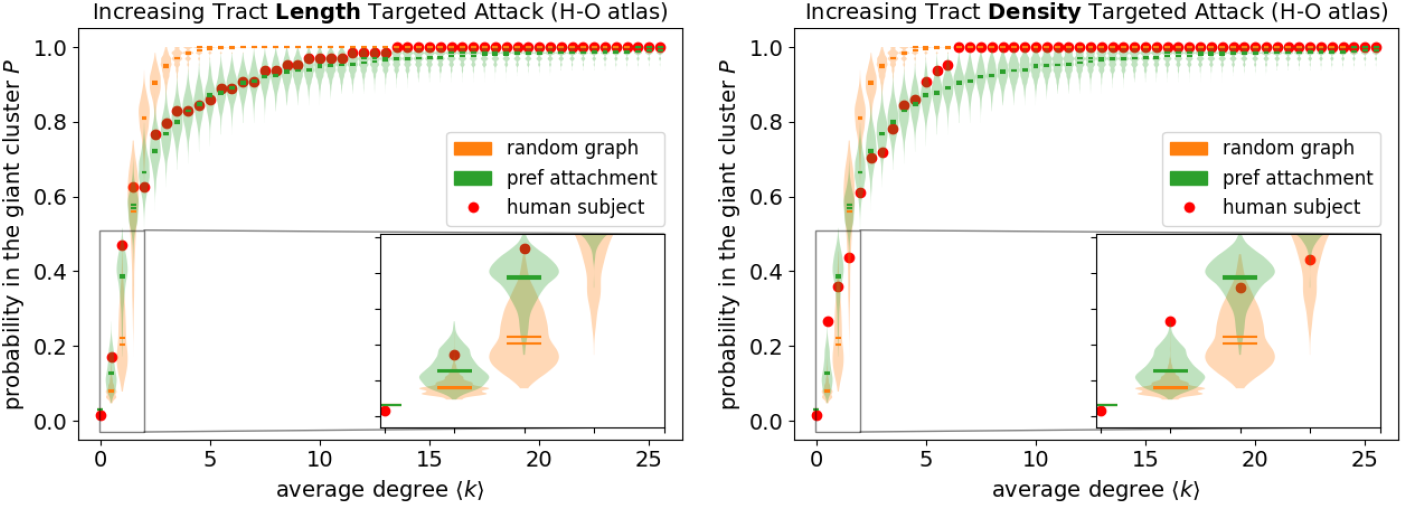
A network built by preferential attachment captures experimental *P* curves at a coarser parcellation, called the Harvard-Oxford (H-O) atlas. Networks are presented as violin plots of simulations (1,000 independent runs). Simulations have the same number of nodes as regions under the H-O atlas (64) as well as the same final average degree of 25.5 seen for the corresponding subject. Experimental curves are derived from dMRI data collected by the UK Biobank under the Harvard-Oxford atlas for one human individual (subject ID: 6025360), the same subject as in Figure 2.

**Figure S6:**
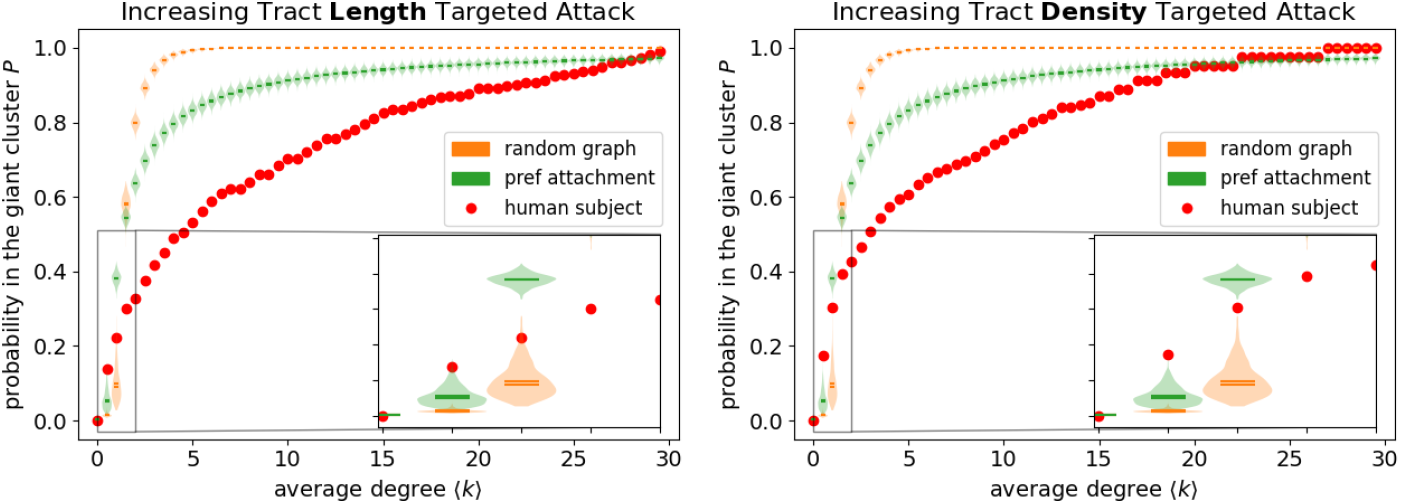
Random and preferential attachment networks are not able to capture *P* curves tabulated with the same number of nodes as regions under the Talairach atlas (727), as well as the same final average degree of 29.9. Violin plots for the networks are determined as in Figure S5. Experimental curves are derived from dMRI data collected by the UK Biobank under the Talairach atlas for one human individual (subject ID: 6025360), the same subject as in Figure 2.

**Figure S7:**
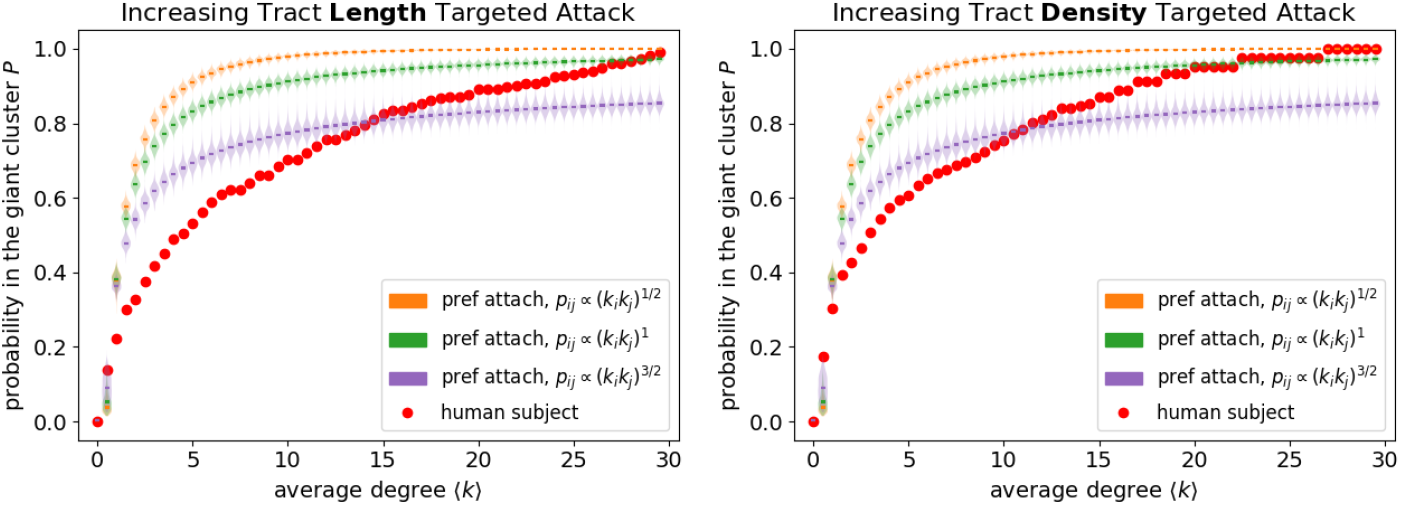
General preferential attachment networks in which the probability of edge formation *p*_*ij*_ is proportional to the multiple of nodes’ *i* and *j* degrees *k*_*i*_ * *k*_*j*_ to some power *x* [63, 64]. What is referred to as preferential attachment in the main text and in Figure S6, corresponds to *p*_*ij*_ ∝ *k*_*i*_*k*_*j*_ in this figure. Despite considering general preferential attachment networks with *x* = 1*/*2 and *x* = 3*/*2, we are still not able to capture *P* curves tabulated with the same number of nodes as regions under the Talairach atlas (727), as well as the same final average degree of 29.9. Violin plots for the networks are determined as in Figure S5. Experimental curves are derived from dMRI data collected by the UK Biobank under the Talairach atlas for one human individual (subject ID: 6025360), the same subject as in Figure 2. Note that we are not able to explore arbitrarily large *x* because those simulation start sampling the same edges over and over, causing the simulation to freeze and not add any new edges.

**Figure S8:**
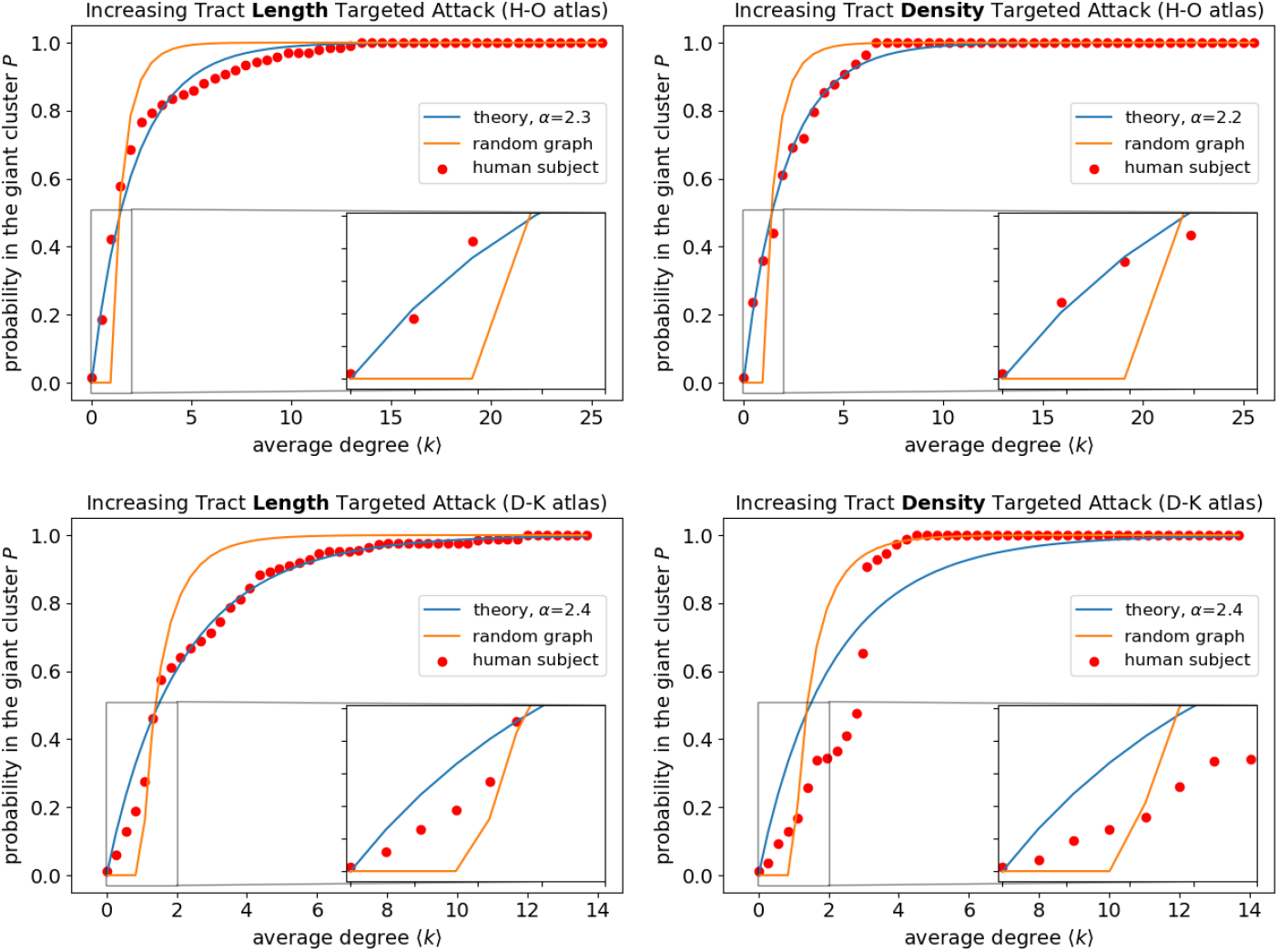
The Giant Cluster Self Preference theory mostly captures *P* curves generated from coarser atlases with smaller fitted *α*. Both atlases considered contain fewer regions than the Talairach atlas (727 regions) used throughout the main text; Harvard-Oxford (H-O) and the modified Desikan-Killiany (D-K) atlases have 64 and 84 regions considered in our analysis, respectively. Curves are derived from dMRI data collected by the UK Biobank for one human individual (subject ID: 6025360), the same subject as in Figure 2. It is not clear why tract density targeted attack under the D-K atlas deviates from the Giant Cluster Self Preference theory. The results look somewhat similar to a random graph and demonstrate a sharp rise in its *P* curve.

**Figure S9:**
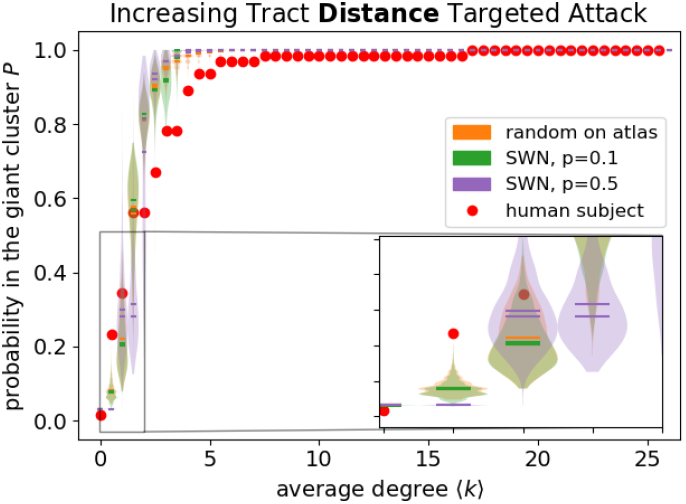
A random network based on actual coordinates of regions according to the Harvard-Oxford (H-O) atlas does not capture *P* curves. Small-world networks (SWNs) are also not able to capture *P* curves tabulated with the same number of nodes as regions under the Harvard-Oxford atlas. Networks are presented as violin plots of simulations (1,000 independent runs). Small-world networks are defined as in [1] and the parameter p in the legend corresponds to the probability of rewiring each edge. For both network types, we first constructed the final network with the correct number of nodes (64) and final average degree (25.5). We then performed targeted attack on increasing tract distance. Note that tract distance is more limited than tract length because it assumes that the trajectory of the white matter tract between the two regions is a straight line. Nonetheless, we find that targeted attack on increasing tract distance provides very similar results to that of tract length (Figure S15). The experimental curve is derived from dMRI data collected by the UK Biobank under the Harvard-Oxford atlas for one human individual (subject ID: 6025360), the same subject as in Figure 2.

**Figure S10:**
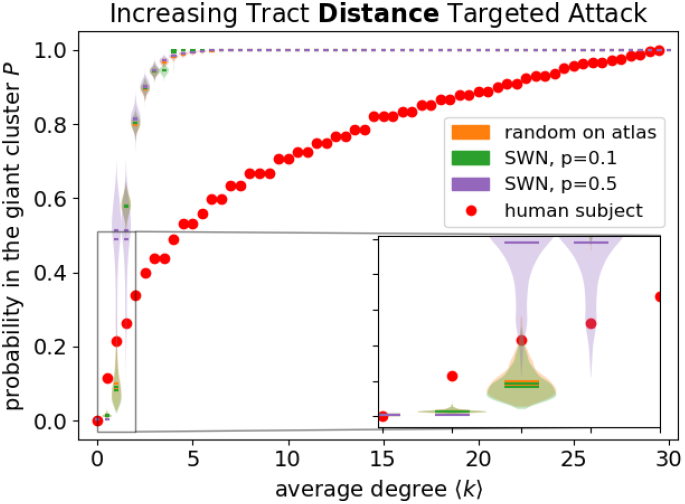
A random graph constrained by the atlas and small-world networks (SWNs) are not able to capture *P* curves tabulated with the same number of nodes as regions under the Talairach atlas (727), as well as the same final average degree of 29.9. Violin plots for the networks are determined as in Figure S9. Curves are derived from dMRI data collected by the UK Biobank under the Talairach atlas for one human individual (subject ID: 6025360), the same subject as in Figure 2.

**Figure S11:**
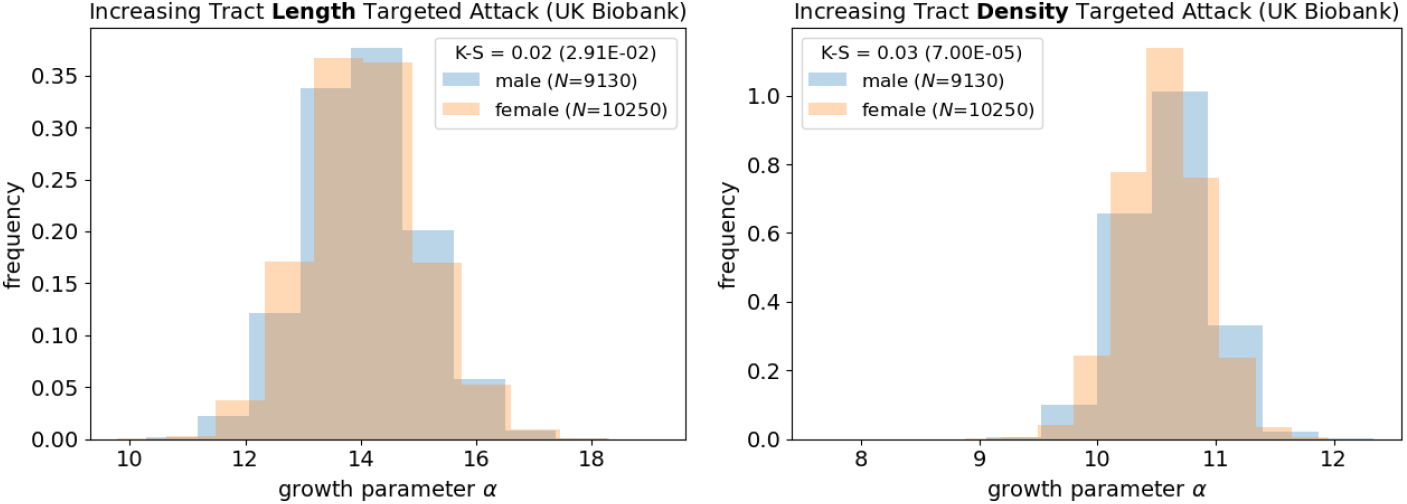
Distributions of the fitted *α* parameter does not vary much between individuals of different sexes. K-S corresponds to the Kolmogorov-Smirnov test statistic, with p-value in parentheses. The variable *N* represents the total number of individuals of the corresponding sex in the UK Biobank.

**Figure S12:**
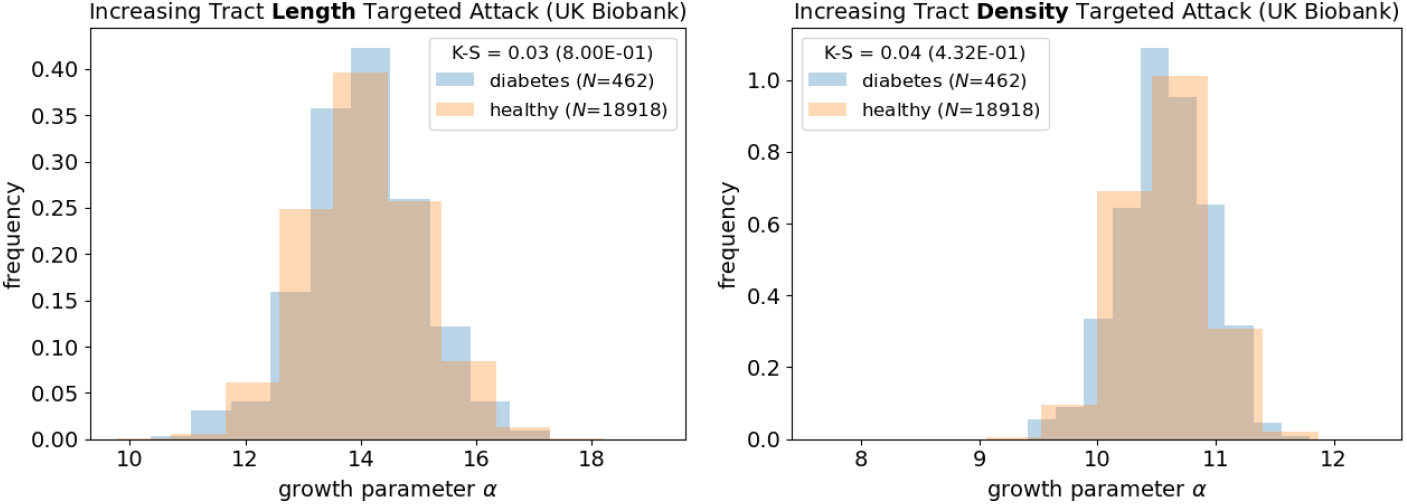
Distributions of the fitted *α* parameter does not systematically vary between individuals diagnosed with diabetes and those not diagnosed with diabetes (healthy). K-S corresponds to the Kolmogorov-Smirnov test statistic, with p-value in parentheses. The variable *N* represents the total number of individuals with the corresponding diabetes diagnosis in the UK Biobank.

**Figure S13:**
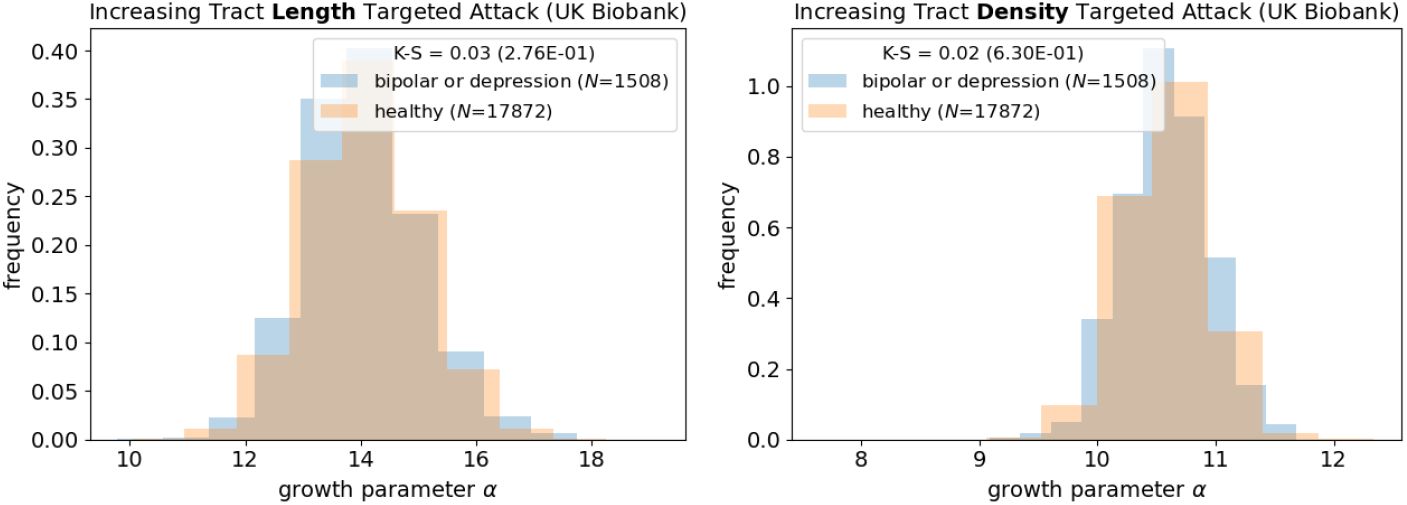
Distributions of the fitted *α* parameter does not systematically vary between individuals diagnosed with bipolar disorder or depression and those not diagnosed (healthy). K-S corresponds to the Kolmogorov-Smirnov test statistic, with p-value in parentheses. The variable *N* represents the total number of individuals with the corresponding bipolar disorder and depression diagnosis in the UK Biobank.

**Figure S14:**
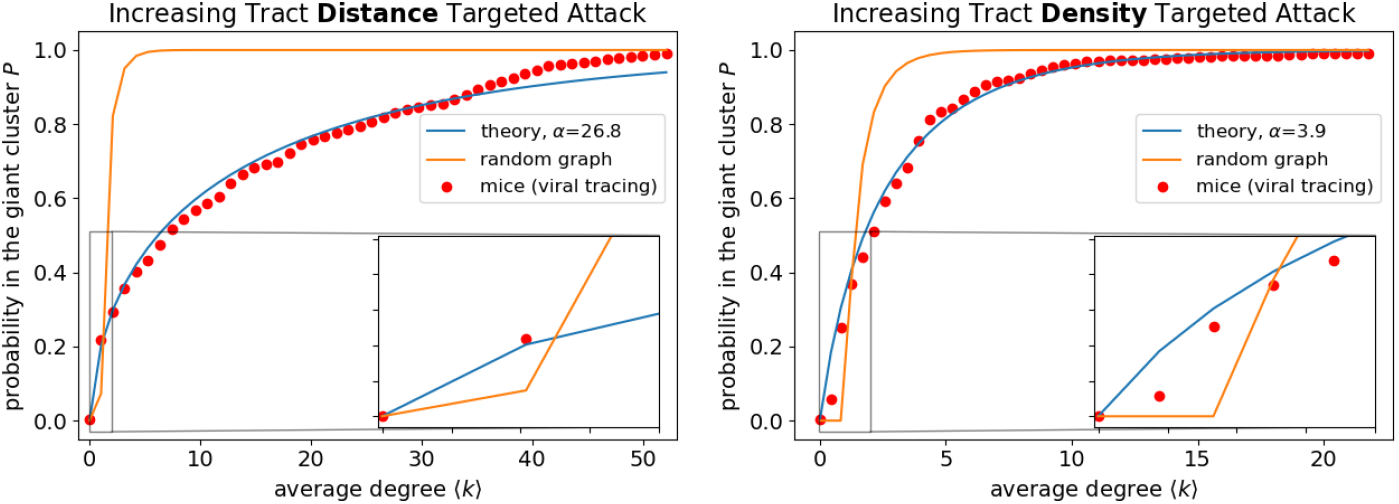
The Giant Cluster Self Preference theory captures the probability for a brain region to be in the giant cluster across average degree with an appropriately fitted *α* parameter for both targeted attack by increasing distance between regions’ center of masses and tract density. Curves are derived from viral tracing data by the Allen Institute averaged over ~ 1000 mice [31]. While *α* values for the two human *P* curves are comparable at 15.3 and 11.0 (Figure 2), the *α* value shown here needed to fit the distance *P* curve is around five times larger than that of the tract density for mouse. The relative differences do not seem to be a result of viral tracing being limited to distances; we find similar trends if we look at corresponding distances for human results (Figure S15). Also, the resolution of the atlas does not seem to be a factor; the atlas for the presented mouse results contains 426 grey matter regions, while that for human contains 727 grey matter regions. Changing the atlas used in processing the dMRI to calculate the connectivity matrix to 84 or 64 regions, yields *α*(distance)/*α*(density) even closer to 1 (Figure S8). Further investigation is needed to elucidate whether relative *α* differences are a result of differences between the dMRI and viral tracing data modalities as highlighted in other studies [65, 66].

**Figure S15:**
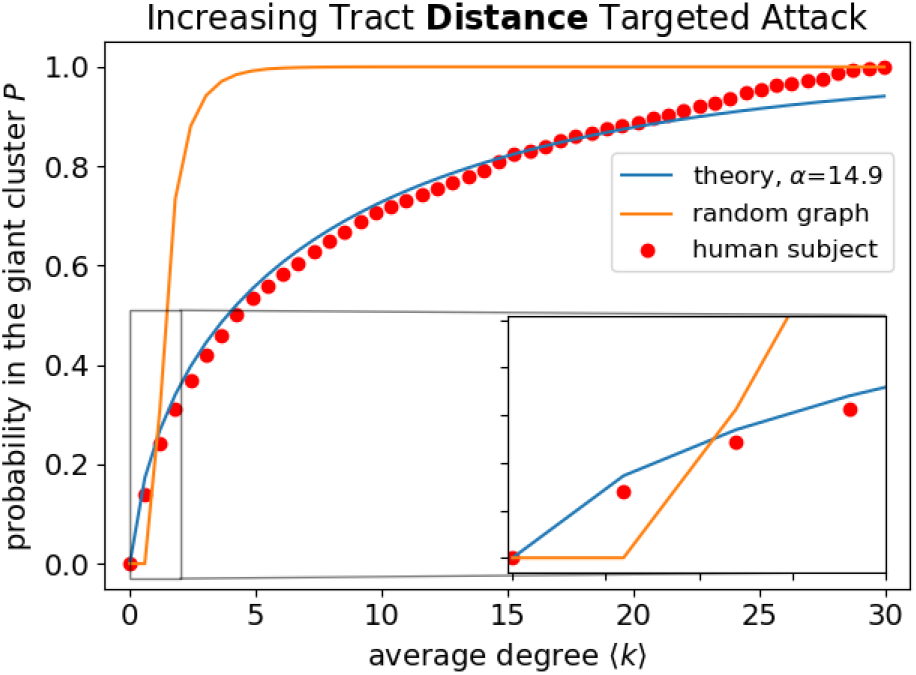
Targeted attack based on center of mass distances of Talairach atlas regions yields similar results as compared to tract lengths. Curves are derived from dMRI data collected by the UK Biobank under the Talairach atlas for one human individual (subject ID: 6025360), the same subject as in Figure 2.

**Figure S16:**
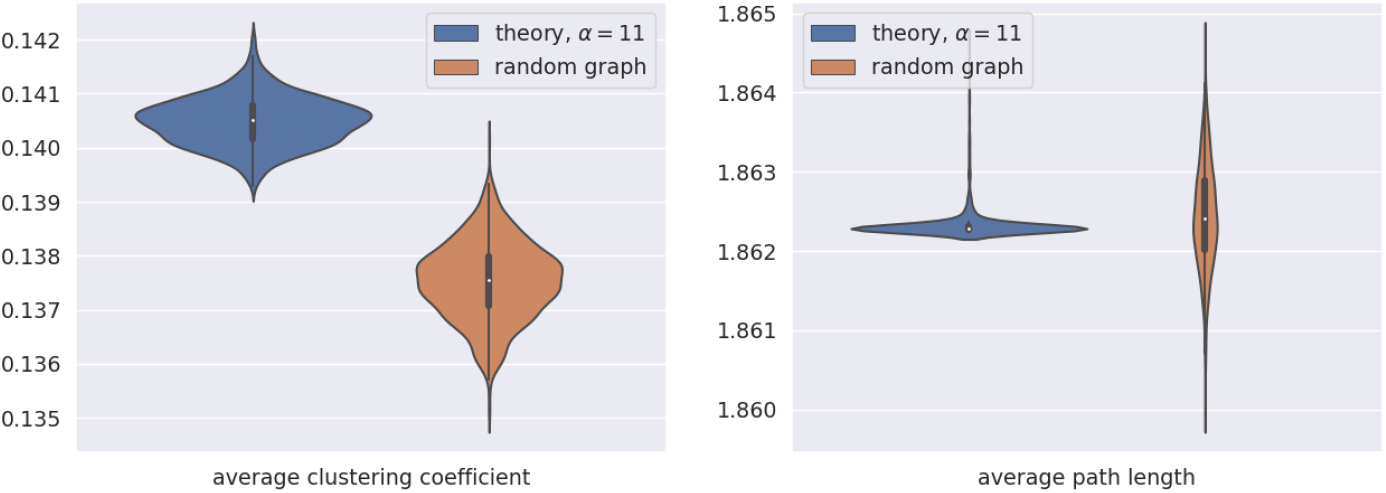
The Giant Cluster Self Preference theory slightly reproduces elevated average clustering coefficients, the hallmark of small-world networks. Our theory’s average clustering coefficient is slightly larger than that of a random graph. In addition, our theory’s average path length is essentially the same as that of a random graph, as predicted by a small-world network. Graph properties are calculated for 1000 independent runs and presented as violin plots with the same number of nodes and final average degree for both graph types. The number of nodes is set to 727, the same number as in the Talairach atlas, and the final average degree is set to 100 to ensure *P* = 1 at the end of each simulation run. Edges are added with *α* = 11 according to the Giant Cluster Self Preference theory.

**Figure S17:**
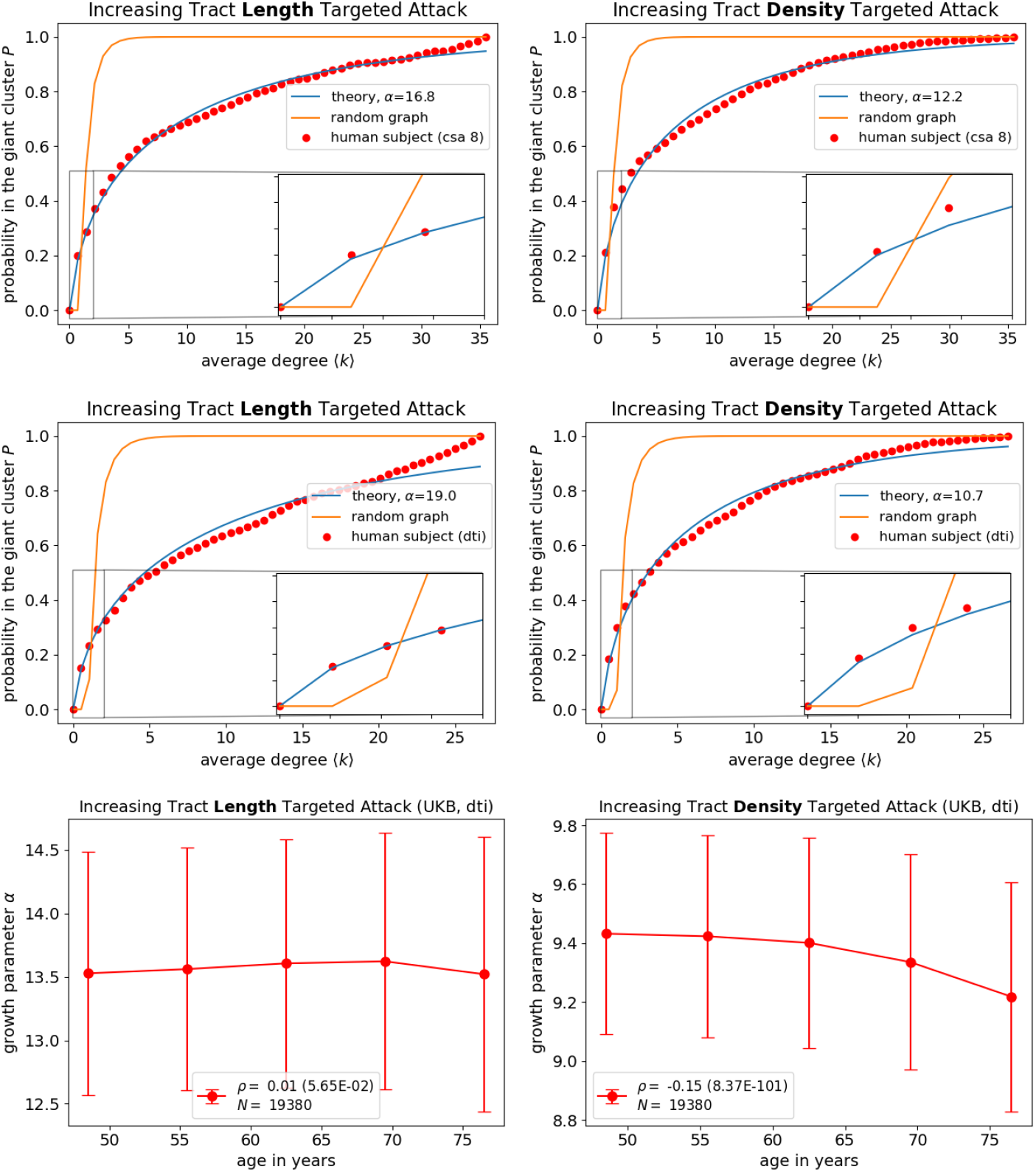
Changing the default procedure in DIPY to calculate the connectivity matrix from dMRI data have little effect on the *P* curves. In the top two figures, we alter the constant solid angle (csa) method to spherical harmonic order 8, compared to order 6 in the main text. In the middle two figures, we use the diffusion tensor imaging (dti) method, rather than the constant solid angle method used in the main text. Curves are derived from dMRI data collected by the UK Biobank under the Talairach atlas for one human individual (subject ID: 6025360), the same subject as in Figure 2. In the bottom two figures, we show results across the entire UK Biobank (UKB) using the dti method. Although increasing tract length targeted attack results are strikingly similar as using the default procedure (Figure 3), those for tract density have slightly lower *α* values that are more correlated with age.

**Figure S18:**
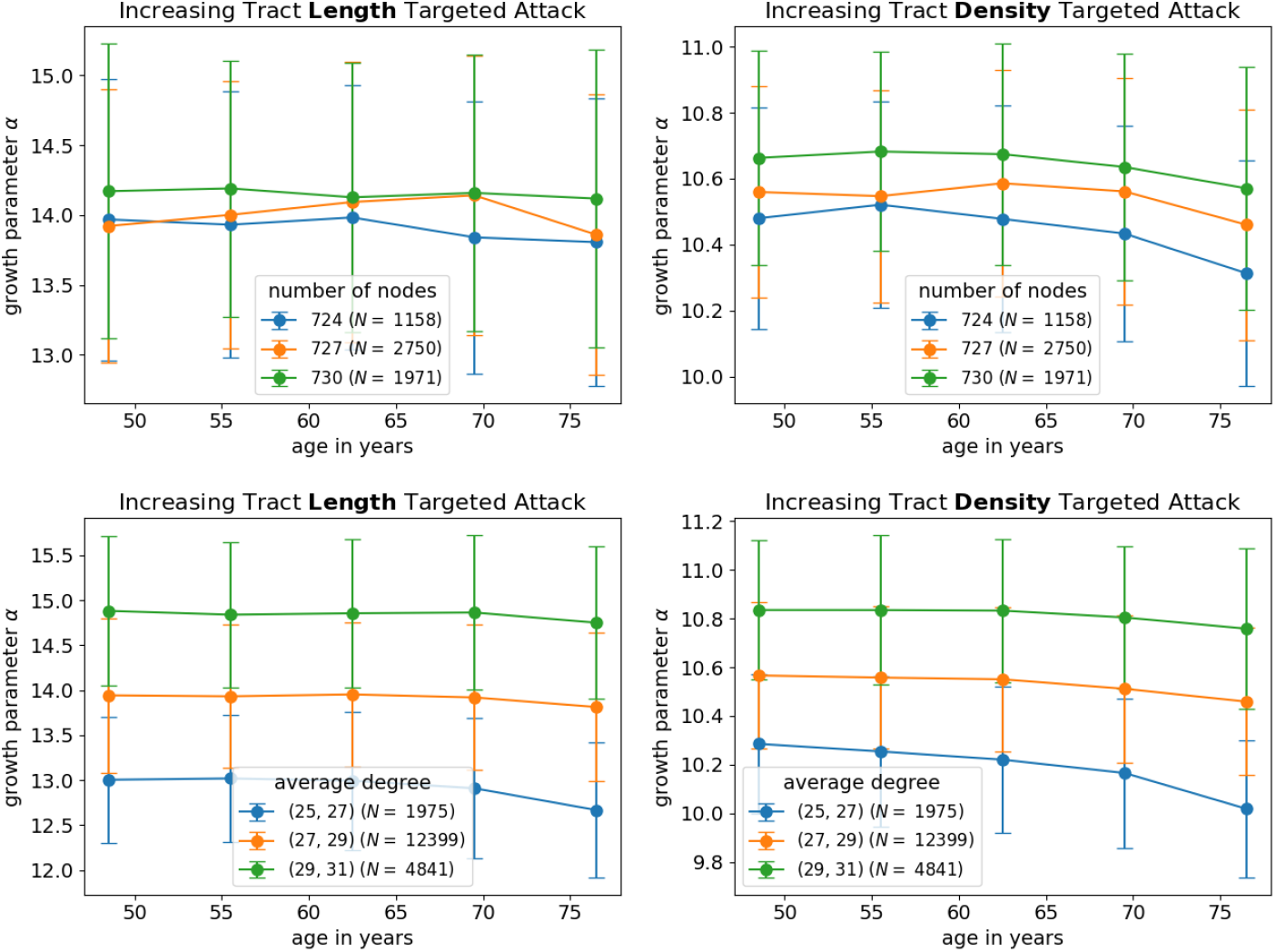
Values of *α* are fairly robust to different number of nodes and average degrees of brain networks in the UK Biobank. Binned data are presented with a line connecting means, and error bars correspond to standard deviations. Thresholds are chosen based on histograms of the respective property (Figure S20). Average degree thresholds encompass a range because values are not discrete. For example (25, 27) corresponds to all networks with average degrees larger than 25 but less than 27. Some variability in *α* is seen as network average degree changes, however, variability is systematic across age such that *α* values differ by a constant.

**Figure S19:**
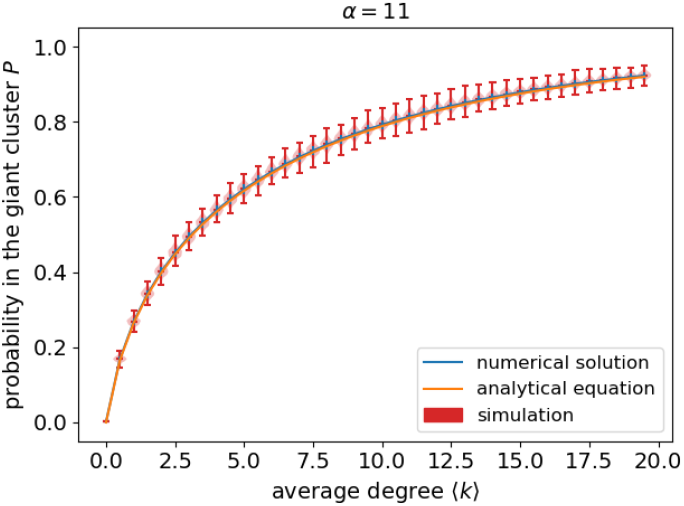
Simulation results verify our theoretical derivation of Equation 9. Violin plots represent 1000 independent runs of a graph with 727 nodes, the same number as in the Talairach atlas, and a final average degree of 100 to ensure that at the conclusion of the simulation, all runs have *P* =1. The probability that an edge is added between nodes in the giant cluster is set to *p* = 100*/*727, while the probability that an edge is added between a giant cluster node and a non-giant cluster node is *p/*11 because *α* is set to 11. The blue line corresponds to the numerical solution of Equation 5, with error bars representing one standard deviation for the respective *p*(*n*|*E*) across *n*. The orange line corresponds to Equation 9 and approximately solves Equation 5 as discussed in the Methods. The analytical equation is simply referred to as ‘theory’ throughout other figures in the main text and SI.

**Figure S20:**
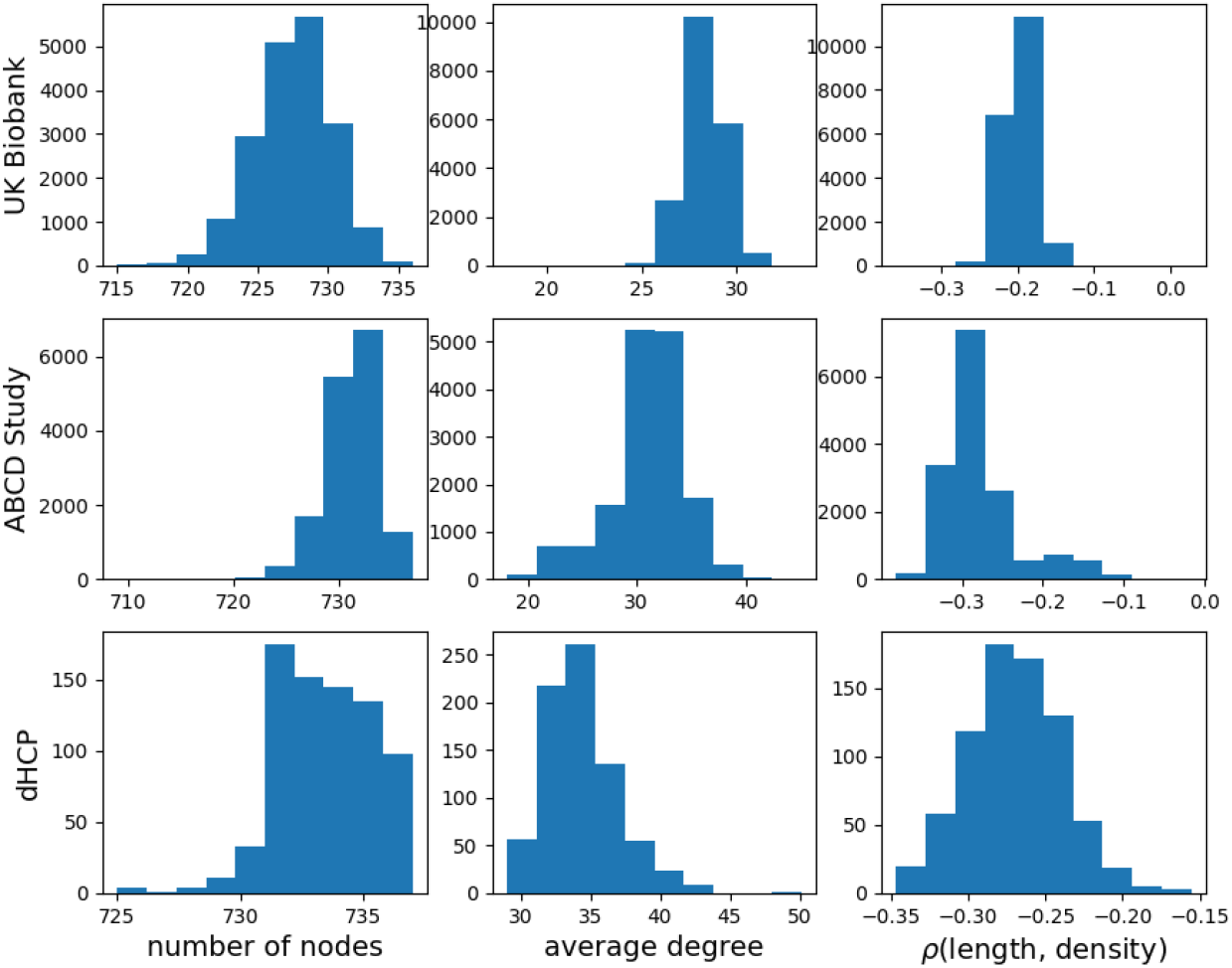
Histograms of basic network properties across individuals in the UK Biobank (*N* = 19380), ABCD Study (*N* = 15593) and dHCP (*N* = 758) data sets. The number of nodes, the average degree of the network before targeted attack is performed, and the Spearman correlation (*ρ*) between tract length and tract density are shown. The y-axis corresponds to the number of individuals with the respective network property value.

## Footnotes

1 Recent brain scans of fetuses in the womb are promising but not yet publicly accessible [11, 12].

2 Limits are known for many graphs such as scale-free and lattices, however the full trajectory from small to large ⟨*k*⟩ remains analytically unsolved [15, 14].

3 Although the Spearman correlation coefficient between *α* and age is highly significant for those *α* values corresponding to tract density with a p-value ~ 10^−19^, the effect size is low with a Spearman correlation coefficient of − 0.06. The Spearman correlation coefficient between *α* and age is not significant for those *α* values corresponding to tract length.

4 These differences in details could be attributed to a combination of factors, such as the particular parcellation of the mouse brain into regions and viral tracing being limited to distances (see Figure S14’s caption). Nonetheless, these results suggest that the *P* curves reflect a generic developmental signal that extends across different species.

5 Notably, results qualitatively differ from ours, in that a critical point in their *P* curves is observed. It is not clear whether the neuronal scale, the *in vitro* sample, and/or the presence of neuronal activity are responsible for differences found compared to our results. Nonetheless, this study showed that *P* = 1 near the expected birth of the fetal mouse [40]. In addition, *P* curves of neurons initially derived from adult mice and fetal mice were similar, indicating that *P* curves reflect a developmental signature that persists throughout organisms’ lifetimes [40, 39].

6 Note that we only need the Giant Cluster Self Preference theory aspect of the Early Path Dominance model to achieve these results since clustering coefficient is a graph theoretical property and edge addition does not depend on growth mechanism in our model.

7 Coarser parcellations yield smaller values of *α* (Figure S8); this makes sense, since all that is needed to connect a new large region into a network is for <u>one</u> of the many sub-regions of that region to get connected, whereas with finer parcellations, it is harder to bring a new region into the network because it is like throwing a dart at a smaller target.

8 Requiring *κ* to remain finite corresponds to an early stage from the growth and development perspective, and to sparsely populated edge spots from the targeted attack perspective. In either case, in the opposite limit - at late stages of growth or when most of the edges are still there in targeted attack, we get the expected result that the connected cluster spans the whole possible network. This approximation is critical to obtaining an analytical expression because it allows us to neglect the *κ* term in the denominator of the transition probability.

9 The average degree ⟨*k*⟩_*g*_ when normalized by the number of nodes present in the growth perspective is equal to *κ/ρ*.

